# Aging seeds of weedy broomrapes and witchweeds lose sensitivity to strigolactones as DNA demethylates

**DOI:** 10.1101/2023.02.27.530112

**Authors:** Guillaume Brun, Jonathan Pöhl, Susann Wicke

## Abstract

Broomrapes (*Phelipanche* and *Orobanche* spp.) and witchweeds (*Striga* and *Alectra* spp.) are obligate root parasitic weeds responsible for major crop yield losses worldwide. Their success in agricultural landscapes is attributable to their ability to produce thousands of long-lived minute seeds that coordinate their germination with the presence of nearby hosts by perceiving host-derived strigolactones. Nevertheless, the processes underlying the alleged decade(s)-long persistence in the field are understudied. Using an accelerated seed aging method coupled to germination and ELISA bioassays, we report that the loss of seed viability and germinability along seed aging is accompanied by a decrease in both strigolactone sensitivity and global DNA methylation. Our results also suggest that seeds of broomrapes are longer-lived than those of witchweeds. Overall, this study deems to initiate further research into how epigenetic mechanisms contribute to alterations in seed viability in parasitic weeds, and how seed aging influence seed responses to their environment.

## INTRODUCTION

A handful of the c. 2000 parasitic plant species of the Orobanchaceae family are weeds, exerting devastating damage on crop yield worldwide. Of particular concern are witchweeds like *Striga hermonthica*, which can cause total cereal crop failures in sub-Saharan Africa, and broomrapes, such as *Phelipanche ramosa*, which feeds of various dicot crops from the west of the Mediterranean basin to the Middle East (Parker 2012, 2013).

Weedy Orobanchaceae, and many of their non-weedy relatives, possess seed traits that render these parasite species highly competitive in agricultural landscapes. First, broomrapes and witchweeds have evolved microspermy, i.e. they produce minute seeds of c. 200 μm in diameter (Murdoch and Kebreab 2013). Second, weedy species show a higher fecundity than their facultative parasitic relatives, allowing them to set thousands of seeds per mature individual in one life cycle (Kebreab and Murdoch 2001, Mourik et al. 2008, Rich and Ejeta 2007, Rodenburg et al. 2006). Third, imbibed seeds undergo a conditioning period of several days to several weeks at 18°C to 30°C (Brun et al. 2019, Lechat et al. 2012, Matusova et al. 2004), before they germinate, mostly strictly in response to picomolar concentrations of host root-derived compounds such as strigolactones (Brun et al. 2018, 2021). Fourth, long-term seed burial experiments and/or predictive modelling indicate that seed longevity in weedy Orobanchaceae (Bebawi et al. 1984, Kebreab and Murdoch 1999, López-Granados and García-Torres 1999, Murdoch and Kunjo 2003) may reach 14−20 years in field conditions, provided sufficient oxygenic conditions and the absence of microbial decay (Gbèhounou et al. 1996, Gbènouhou 1998, Mourik et al. 2005, Murdoch and Kebreab 2013). Therefore, the production of many easily dispersed, long-lived seeds that coordinate their germination with the presence of a host largely compensates for a low attachment success (López-Granados and García-Torres 1993) and hampers parasite management efforts.

Molecular and physiological changes associated with seed aging are understudied in parasitic plants, although such research would greatly improve our understanding of long-term persistence and facilitate the design of integrative weed control strategies. Early studies in non-parasitic plants indicate that breaks in DNA, epigenetic variations, and abnormal gene expression patterns accumulate during seed aging, thereby increasing the frequency of affected cell damage that eventually exceeds the threshold beyond which seed germinability decreases (Zhang et al. 2021). The kinetics of germination-affecting changes in parasitic plants as their seeds age remain unknown.

Here, we aimed to understand the consequences of seed aging in weedy broomrapes and witchweeds. First, we evaluated how the germination dependence for host-derived compounds and sensitivity towards those compounds change in aging seeds of selected Orobanchaceae weeds as viability declines. Second, we explored whether seed aging in weedy parasites associates with modifications of the epigenetic landscape. To this end, we recorded seed viability, sensitivity to germination stimulants, and global DNA methylation levels over time in the context of an accelerated aging method.

## MATERIAL & METHODS

### Plant material

Seeds of *Phelipanche ramosa* (L.) Pomel, growing on winter oilseed rape in 2017 in Saint-Maxire (Deux-Sèvres, France), were kindly provided by the Delavault lab at the University of Nantes (France). Seeds of *Striga hermonthica* (Del.) Benth., collected in 2022 in a sorghum field in the Kibos district (Kisumu, Kenya), and *Alectra vogelii* (Del.) Benth., growing in a cowpea field in the Embu district (Embu, Kenya) were kindly provided by the Runo lab at Kenyatta University, Nairobi (Kenya). Seeds of *Orobanche crenata* Forssk. collected in a carrot field near Ramat David, Israel, in 2019 were kindly provided by Radi Aly (Agricultural Research Organization, Newe Yaar Research Center). Seeds of *Aeginetia indica* (L.) Huth were harvested in 2022 from a permanent living collection of the parasite growing on *Setaria geniculata* at the Späth-Arboretum of HU Berlin (Germany). All seeds were filtered through a sieve to isolate individuals of 200−224 μm in diameter; cleaned seeds were stored in the dark at room temperature until use.

### Accelerated seed aging

Plastic containers (2.5 − 5 L capacity) were filled with 100 - 200 mL saturated sodium nitrite solution (NaNO_2_), which we prepared by dissolving crystals in deionized water. The containers were incubated at 50 °C for a day to reach a relative humidity equilibrium of 60 %. Dry seeds were spread as a thin layer onto watch glasses, inserted into the plastic containers at 50°C. We collected seeds in regular intervals until 240 h of incubation. Collected seeds were allowed to re-adjust to room temperature and relative humidity for an hour before proceeding to surface-sterilization.

### Seed surface-sterilization

Seeds were immersed in bleach, diluted to 4 % chlorine with deionized water, for five minutes, followed by three washing steps of 30 s each and three washing steps of five minutes each in autoclaved water (Brun et al. 2019 for details). For germination and viability assays, 10−20 mg surface-sterilized seeds were resuspended at a 10 mg.ml^−1^ concentration in an incubation medium consisting of 0.1% Plant Preservative Mixture (PPM, Plant Cell Technology, USA) and 1 mM HEPES buffer, adjusted to pH 7.5 with KOH. Fifty microliters of seeds were distributed into 96-well plates (Greiner Bio-One) and the volume was adjusted to 90 μl with incubation medium. For DNA extraction, 30 mg surface-sterilized seeds were resuspended in 10 mL incubation medium, and the suspension was transferred into cell culture flasks. 96-well plates were sealed with parafilm, and plates and flasks were wrapped in aluminum foil before their transfer to species-optimal seed conditioning environments. That is, seeds of *P. ramosa* and *O. crenata* were incubated for seven days at 21 °C. Seeds of *S. hermonthica* and *Al. vogelii* were incubated for eight days at 31 °C. Seeds of *Ae. indica* were incubated for twelve days at 28 °C.

### Germination and seed viability assays

For germination assays, conditioned seeds of *P. ramosa, O. crenata, S. hermonthica*, and *Al. vogelii* were stimulated with 10 μl of 10^−5^ to 10^−12^ M of a racemic mixture of the strigolactone analog GR24 (hereafter referred to as *rac*-GR24), diluted in 0.1% acetonitrile. Imbibed seeds of *Ae. indica* were stimulated with 10 μl of various concentrations of coconut water (10^0^ X to 10^−7^ X). Seeds stimulated with 10 μl acetonitrile 0.1% or water, respectively, were used as negative controls. After 4 days of incubation, seed germination was assessed under a SMZ-168 microscope, connected to a Moticam S6 camera. Seeds were considered germinated when the radicle protruded out of the seed coat. For viability assays, imbibed seeds were transferred to 100 μl of 1% 2,3,5-tryphenyl tetrazolium chloride (TTC) for six days at 21−31 °C, depending on the species-specific temperature optimum. Except for *Al. vogelii* seeds, seeds were further incubated in bleach containing 4 % chlorine for five minutes (*P. ramosa* and *O. crenata*), 3% chlorine for three minutes (*Ae. indica*), or 4 % chlorine for three minutes (*S. hermonthica*) in order to unequivocally identify positively stained embryos. We considered seeds as viable when a homogenous red staining within the embryo showed. Each experiment was replicated at least three independent times.

### Global DNA methylation quantification

Imbibed seeds from three biological replicates were blotted on tissue paper and snap-frozen in liquid nitrogen. Genomic DNA was extracted using the DNeasy Plant Pro Kit (Qiagen), with minor adjustments to the manufacturer’s instructions. That is, seeds were first grinded to a fine powder in liquid nitrogen before resuspending the ground material in 500 μl of buffer CD1. The suspension was then transferred to tissue disruption tubes, which were vortexed for 10 minutes at room temperature. Genomic DNA was eluted in 50 μl of EB buffer. We measured the DNA concentration using a Qubit4 Fluorometer in 1X dsDNA High Sensitivity mode. Relative quantification of 5-methylcytosine (5-mC) over time was determined using the 5-mC DNA ELISA Kit (Zymo Research), using 100 ng of genomic DNA per sample as input. All biological replicates were assayed in duplicates. Absorbance at 405 nm was measured after 30 minutes, using a Multiskan SkyHigh microplate reader. Variations in 5-mC content over accelerated aging were expressed relatively to the absorbances at 405 nm in non-aged samples.

### Statistical analyses

All statistical analyses were carried out using R version 4.2.2 (R Core Team, 2022). Dose and time-response curves were analysed using the ‘drc’ package (Ritz et al. 2015). First, a three-parameter log-logistic model was fitted for each dataset. Second, we selected the best model among linear, quadratic, and cubic regression models using the mselect function. The model displaying the smallest Akaike’s Information Criterion (AIC) and residual standard error, as well as the larger p-value from the lack-of-fit test against one-way ANOVA model was selected as the best model to fit the data. The ED function was then applied to compute longevity estimates expressed as the time necessary to reach 5, 50, and 95% loss of germination or viability, and sensitivity estimates expressed as the concentration of chemicals to observe 50 % of the maximum germination.

## RESULTS & DISCUSSION

### Weedy broomrapes are longer-lived than witchweeds

We assessed the longevity of the strigolactone-insensitive parasitic weed *Ae. indica* and four strigolactone-sensitive weedy Orobanchaceae (*O. crenata, P. ramosa, Al. vogelii, S. hermonthica*) through an accelerated seed aging experiment. Seed viability, expressed as the percentage of seeds displaying positive TTC staining, was overall significantly higher than germination for all species except for *Ae. indica* (Fig. 1A). This result was expected, as fully or close to fully viable seed batches are rarely fully competent for germination, even under optimal germination conditions (Gbèhounou et al. 2003, Kebreab and Murdoch 1999, Thorogood et al. 2009). However, we noted that the difference between viability and germination increased as seeds aged. For instance, in *P. ramosa, O. crenata*, and *S. hermonthica*, we observed 96.9%, 96.12%, and 82.3%, of viable non-aged seeds germinated, respectively, as opposed to 68.3%, 72.6%, and 20.5% after 48 hours of accelerated aging (Fig. 1A). Three-parameter Weibull type 1 models, which we used to fit survival curves and compute longevity estimates, showed that seeds of *S. hermonthica* and *Ae. indica* were significantly faster than the other species to lose 5% germinability (1.2 ± 0.4 h and 4.9 ± 0.7 h, respectively). In *Al. vogelii*, the initial viability of 35.1 ± 3.7% was significantly lower than that of any other species, which may explain why it lost 5 % germinability slower than other species, even though it was the fastest to lose 5% viability (4.7 ± 2.1 h). We observed that seeds of the broomrape species *O. crenata* seeds took 74.6 ± 3.5 h and 272.2 ± 18.2 h to lose 50 % and 95 % viability, respectively, as opposed to 64.9 ± 2.9 h and 202.9 ± 6.5 h for *P. ramosa* (Fig. 1B). This corroborates a previous prediction, based on various saturated salt environments, that *O. crenata* seeds are slightly longer-lived than those of *P. aegyptiaca*, a closely related sister species of *P. ramosa* (Kebreab and Murdoch 1999). Most importantly, we found that the broomrape species are more resistant to aging and significantly longer-lived than witchweeds, which took 121.3 ± 15.7 h (*Al. vogelii*), 100.7 ± 5.7 h (*S. hermonthica*), and 48.4 ± 2.8 h (*Ae. indica*) to lose 95% viability (Fig. 1B).

**Fig. 1.**
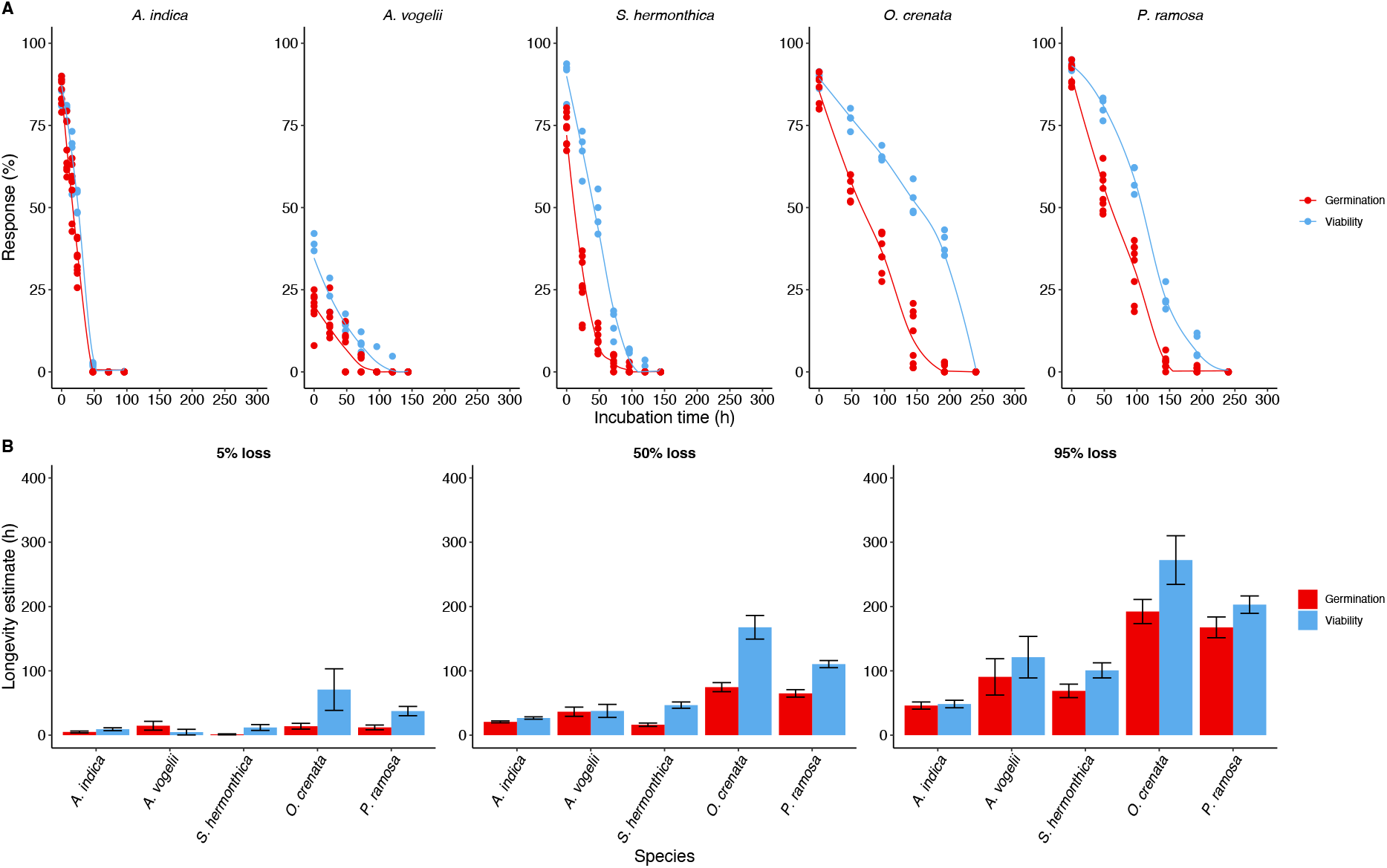
Seed longevity of broomrapes and witchweeds under an accelerated aging protocol. **A**. Seeds incubated for 0-240 hours in a saturated sodium nitrite environment (50°C, 60% relative humidity) were conditioned and treated with 100% coconut water (*A. indica*) or 10−6 M *rac*-GR24 (*A. vogelii, S. hermonthica, O. crenata, P. ramosa*). Germination was recorded after 4 days (n = 8). Viability was measured by incubating conditioned seeds in 1% TTC for 6 days (n = 4). **B**. Longevity estimates expressed as 5%, 50%, 95% loss of viability or germinability according to a three-parameters Weibull type 1 model. Error bars indicate confidence intervals at a 95% confidence level.

No robust sources estimating the longevity of *Al. vogelii* or *Ae. indica* seeds are available. However, unsourced data reported in several instances that *S. hermonthica* seeds would survive in the soil for more than ten years. Yet, conflicting literature reports indicated 52 % depletion of the soil seed bank over two wet seasons in Gambia (Murdoch and Kunjo 2003) and 75 % loss of viability of *S. hermonthica* seeds over six months of burial in Benin (Gbèhounou et al. 1996) and up to total loss over two wet seasons (Gbènouhou 1998). Such steeper decrease may, however, have been overestimated due to the use of small mesh packets of unrealistically high seed density that may have triggered higher microbial decay (Mourik et al. 2005). Seeds of *O. crenata* buried in the soil in Spain for one and twelve years lost their viability to 30 % and 98.7 %, respectively (López-Granados and García-Torres 1999). This seemingly matches a predicted 38.5 % loss of viability of *P. aegyptiaca* seeds over a full year in an Eritrean unimodal rainfall area (Kebreab and Murdoch 1999). Similarly, 10 % of *Striga asiatica* seeds remained viable after 14 years of burial at 152 cm depth (Bebawi et al. 1984). However, most of the witchweed seed population accumulates in upper soil layers (Robinson and Kust 1962), where the total loss of viability occurs between five and nine years (Bebawi et al. 1984).

We argue that, contrary to past assumptions, broomrape seeds might be longer-lived than those of witchweeds. Although our study did not cover a taxonomically representative array of species from each clade of Orobanchaceae, we report that one-year-old batches of witchweed seeds survive significantly less long than four to six-year-old batches of broomrape seeds in the context of an accelerated aging experiment. Longevity is typically multi-factorial and depends notably on the initial viability of seed batches. Therefore, it is unsurprising that *Al. vogelii* displayed a short life-span in our accelerated aging experiment, given its low viability compared to the other species (Fig. 1). We are not the first to report that germination of *Al. vogelii* seeds is low (Mallu et al. 2022, Okonkwo and Nwoke 1975), which may reflect a naturally low seed viability for this species. Despite the fact that *S. hermonthica* has a far greater initial viability than *Ae. vogelii*, we report similar longevity estimates for both species. This leads us to hypothesize that at equally high initial viability, *Al. vogelii* seeds are naturally longer-lived than *S. hermonthica* seeds.

From an anatomical viewpoint, the reduction in seed size often associates with a reduction of seed cellular complexity along the transition to obligate parasitism. All species presented here have reduced endospermic tissue compared to many facultative hemiparasites and nonparasites (Joel and Bar 2013), although notable exceptions exist in genera like *Lathraea* spp., *Conopholis* spp., or *Cistanche* spp. Meta-analyses of seed traits have recently indicated that non-endospermic seeds are on average 3.3-fold longer-lived than endospermic seeds (Merritt et al. 2014, Probert et al. 2009, Tausch et al. 2019). It will be necessary to test whether endosperm structure, cellular and/or metabolic composition, and functionality between seeds of these species explain the apparent differences in longevity. Finally, the obligate parasites tested here belong to distinct clades of Orobanchaceae (Schneeweiss 2013), which represent distinct trophic specializations and diverse seed traits. For example, while *Orobanche* spp. and *Phelipanche* spp. belong to a group of germination stimulant-sensitive holoparasites within Orobanchaceae, *S. hermonthica* is phylogenetic closely related to *Buchnera* spp., the latter of which containing several facultative hemiparasites with larger, strigolactone-insensitive seeds (Krause and Weber 1990, Okonkwo and Nwoke 1974). The same holds true for the hemiparasites *Escobedia* spp., which are close relatives of *Alectra* spp. (Cardona-Medina and Ruiz 2015). The most parsimonious explanation would, therefore, be that the transition to obligate parasitism is the result of a convergent evolution within the Orobanchaceae, which implies independent acquisitions of strigolactone dependency and longevity traits.

### Chemical germination requirements narrow as seeds of strigolactone-sensitive species age

We assessed whether chemical requirements for seed germination vary over periods of aging. To this end, we tested decreasing concentrations of *rac*-GR24 on conditioned seeds, which were subjected to accelerated aging for 0 to 240 h (Fig. 2). A maximum concentration of 10^− −6^ M was consistently found to be most potent for germination regardless of seed age. Fitting three-parameters log-logistic models to model dose-response curves and compute the sensitivity to *rac*-GR24 expressed as the concentration of molecule at which 50% of the maximum germination is observed (EC_50_, Table 1) revealed that non-aged seeds of *P. ramosa* were ca. 100-fold more sensitive to *rac*-GR24 than all other species. This results was expected given that batches of *P. ramosa* seeds are often sensitive to synthetic or natural strigolactones in the picomolar range, while the sensitivity of *Striga* and *Orobanche* species rarely drop below the nanomolar range (Ćavar et al. 2015). Interestingly, strigolactone sensitivity decreased as seed aged, the gradation of which differed between species (Table 1). For instance, strigolactone sensitivity decreased by about 100-fold every 48h of accelerated aging in *P. ramosa*, indicating that they were about 10000-fold less sensitive to *rac*-GR24 by the time they lost 50% viability (Figure 1B, Table 1). We also found that the sensitivity to *rac-*GR24 was about 10-fold lower in 8 and 9-year-old batches of *P. ramosa*, in correlation with significantly lower maximum germination rates of 72.0 ± 3.0% and 59.2 ± 2.8%, respectively (Fig. 3). Similarly, *O. crenata* and *Al. vogelii* seeds were found 1000-fold and 10000-fold less sensitive to *rac*-GR24 after 144h and 24h, respectively (Table 1), which correlates viability losses of more than 50% (167.7 ± 8.8 h and 37.7 ± 4.9 h, respectively; Fig. 1B). A similar trend was found for *S. hermonthica*, although it took 72h to observe a significant decline of *rac*-GR24 sensitivity by 10-100-fold (Table 1) while 50% viability loss was reached after 46.7 ± 2.4 h only (Fig. 1B). Interestingly, 6-year-old seeds of *S. hermonthica* displayed maximum germination of 22.9 ± 3.7% germination and we estimated its sensitivity to *rac-*GR24 to be 5000-fold less than that of 1-year-old seeds (Fig. 3).

**Fig. 2.**
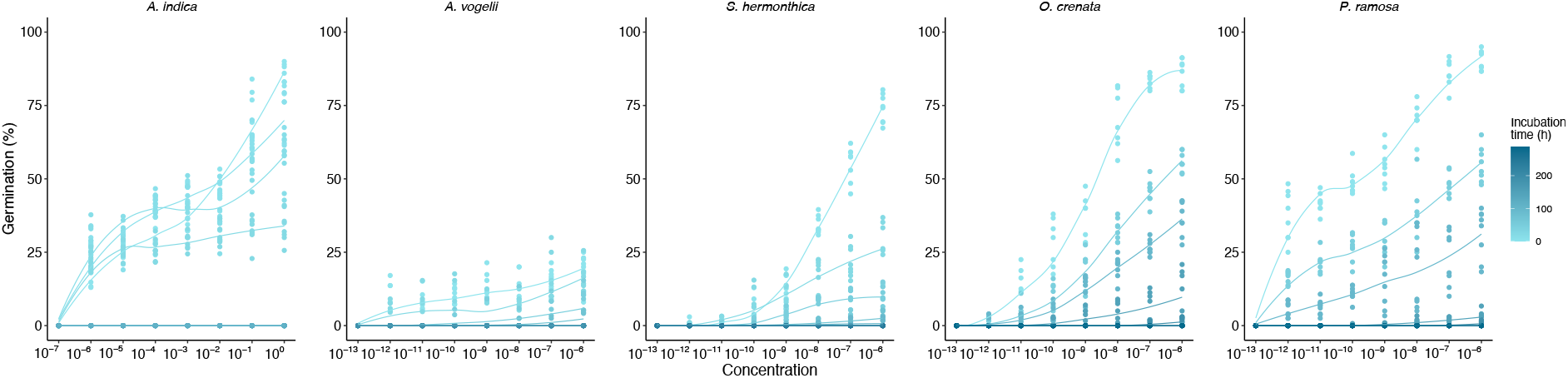
Accelerated seed aging influences seed sensitivity to germination stimulants. Seeds incubated for 0−240 hours in a saturated sodium nitrite environment (50°C, 60% relative humidity) were conditioned and treated with decreasing concentrations of coconut water (*A. indica*) or *rac*-GR24 (*A. vogelii, S. hermonthica, O. crenata, P. ramosa*). Germination was recorded after 4 days (n = 8).

**Table 1.** Sensitivities to germination stimulants after accelerated aging (EC_50_ ± SE)

|  | 0h | 8h | 16h | 24h | 48h | 72h | 96h | 120h | 144h |
| --- | --- | --- | --- | --- | --- | --- | --- | --- | --- |
| *A. indica* | $0.6 \pm 1.1 \times 10^{-1}$ | $2.8 \pm 4.5 \times 10^{-5}$ | $1.5 \pm 0.6 \times 10^{-6}$ | $6.2 \pm 1.1 \times 10^{-6}$ | nd. | nd. | nd. | nd. | nd. |
| *A. vogelii* | $0.9 \pm 6.2 \times 10^{-8}$ | nd. | nd. | $1.6 \pm 1.3 \times 10^{-4}$ | $0.5 \pm 7.5 \times 10^{-4}$ | $1.3 \pm 1.5 \times 10^{-3}$ | nd. | nd. | nd. |
| *S. hermonthica* | $0.3 \pm 6.3 \times 10^{-9}$ | nd. | nd. | $3.8 \pm 3.1 \times 10^{-9}$ | $0.2 \pm 6.3 \times 10^{-9}$ | $2.3 \pm 7.6 \times 10^{-7}$ | nd. | nd. | nd. |
| *O. crenata* | $1.4 \pm 0.4 \times 10^{-9}$ | nd. | nd. | nd. | $9.4 \pm 3.4 \times 10^{-9}$ | nd. | $2.4 \pm 1.9 \times 10^{-8}$ | nd. | $0.4 \pm 2.8 \times 10^{-6}$ |
| *P. ramosa* | $2.0 \pm 3.6 \times 10^{-11}$ | nd. | nd. | nd. | $1.0 \pm 1.9 \times 10^{-9}$ | nd. | $1.2 \pm 5.2 \times 10^{-7}$ | nd. | $0.3 \pm 1.3 \times 10^{-6}$ |

**Fig. 3.**
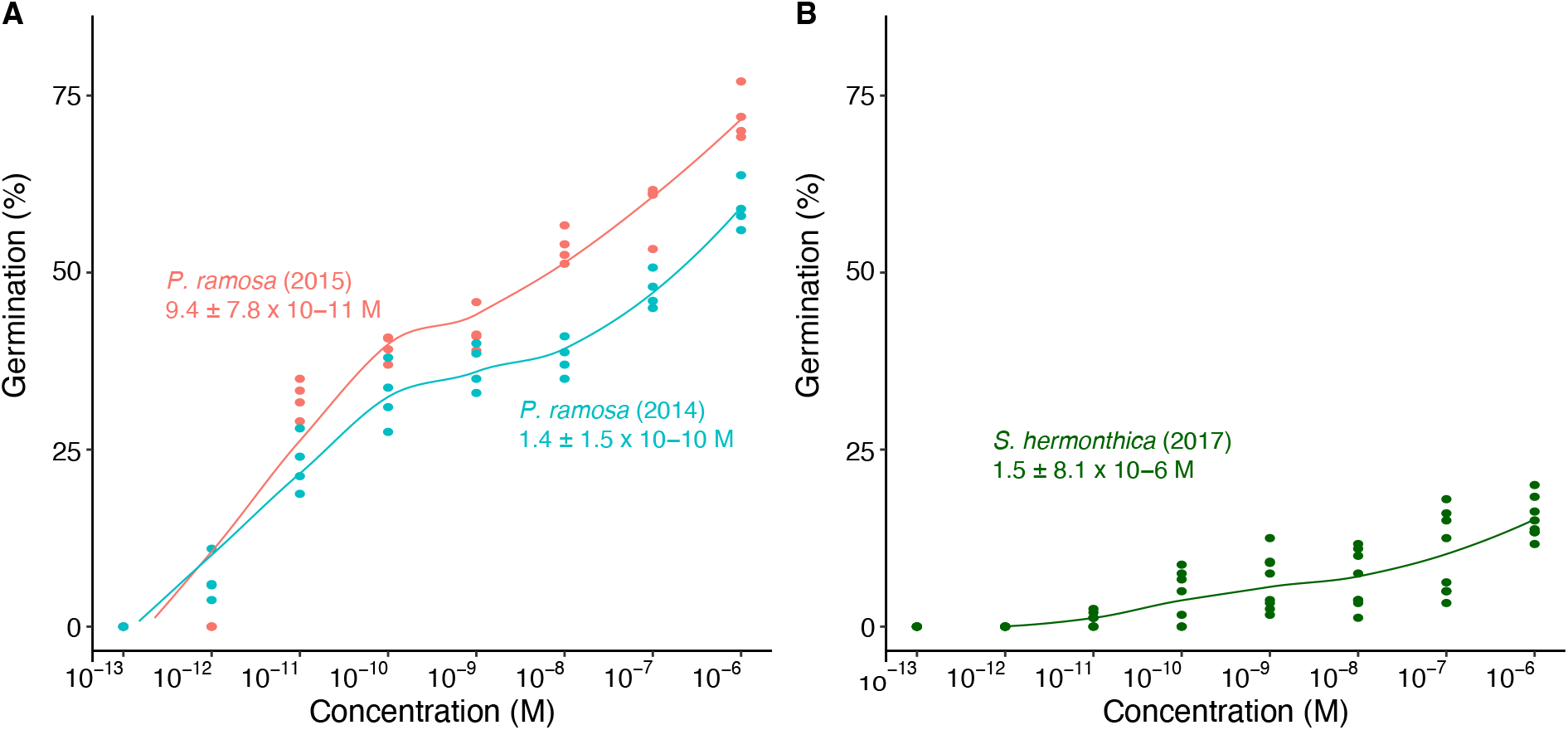
Reduced germinability and sensitivity to strigolactones in naturally aged populations of *P. ramosa* and *S. hermonthica*. **A**. Germination of *Phelipanche ramosa* seeds harvested in a winter oilseed rape field in 2014 (blue) and 2015 (red) upon decreasing concentrations of *rac*-GR24 (n = 4). **B**. Germination of *S. hermonthica* seeds harvested in a sorghum field in 2017 upon decreasing concentrations of *rac*-GR24 (n = 8). Sensitivity to rac-GR24 according to a three-parameter log-logistic model are indicated (EC_50_ ± SE).

Our data indicate that seed aging significantly constraints seed responses to host-derived stimulants. To germinate, older seeds must perceive much higher concentrations of strigolactones. An exception from this observation is the strigolactone-insensitive *Ae. indica*, whose germination in high concentrations of coconut water decreased as seed aged, while germination rates remained stable at concentrations ranging from 10^−4^ to 10^−6^ % (Fig. 3). As a result, sensitivity to coconut water increased by 10 000-fold within 24 h, at which time a 50% viability loss occurred (Table 1; Fig. 1B). Coconut water has also been reported in several instances as stimulating germination of nonparasitic species such as *Cucumis sativus* (Cucurbitaceae, Oka 2014), *Dracontium grayumianum* (Araceae, Torres et al. 2011), *Veitchia merilli* (Arecaceae, Trisnaningsih and Wahyuni 2020), and *Allium ascalonicum* (Amaryllidaceae, Sudaryono and Prahardini 2021). Coconut water contains notably quantities of germination-inducing phytohormones like auxin, cytokinins, and gibberellins (Yong et al. 2009), all of which known to stimulate germination of *Ae. indica* seeds in a dose-dependent manner (Kato and Hisano 1983). Root exudates of many nonparasitic species contain cytokinins and auxin (e.g. Reddy et al. 1989, Eichmann et al. 2021), suggesting that coconut water mimicked the effect of root exudates in stimulating germination of *Ae. indica*. Seeds of *Ae. indica* would become more sensitive to nearby root exudates as they age, thereby increasing the probability to germinate even at low(er) host (root) density. This contrasts the adaptations of strigolactone-requiring species, whose probability of germination decreases not only as a function of viability but also at reduced host density. Further research with improved taxonomic representation is needed to test whether this aging-associated germination response is a specific adaptation to agricultural environments or reflects ecological preferences to a tropical habitat (*Ae*. indica) rather than subtropical or semi-arid habitats of witchweeds and broomrapes, respectively.

### Global DNA demethylation occurs during aging of strigolactone-requiring species

Viability loss along seed aging coincides with decreasing global DNA methylation, regardless of seed storage behavior (Michalak et al. 2013, 2015a, 2015b, 2022, 2023, Plitta-Michalak et al. 2018, 2021, 2022). That is why, we assessed whether changes in the DNA methylation status occurred during accelerated aging of all parasitic species. Witchweed seeds were collected after 24 and 48 h of incubation, whereas longer-lived broomrape seeds were collected after 72 and 120 h of incubation in the saturated sodium nitrite environment. DNA was extracted for non-aged and aged seeds conditioned for seven days in order to quantify changes in 5-mC content relative to the initial abundance of 5-mC motifs in non-aged samples (Fig. 4). We observed a significant decline in global DNA methylation for all strigolactone-requiring species. A higher extent of DNA demethylation was found in *Al. vogelii* with 16.7 to 23.2% loss in 5-mC content after 24 and 48h of accelerated aging. Changes in 5-mC content were subtler in the three other species, with a maximum decline of 5.9 %, 4.1 %, and 7.6 % at the final aging time point in *S. hermonthica, O. crenata*, and *P. ramosa*, respectively. We observed no notable demethylation in *Ae. indica* after 24 h of incubation; measurements for 48 h samples failed.

**Fig. 4.**
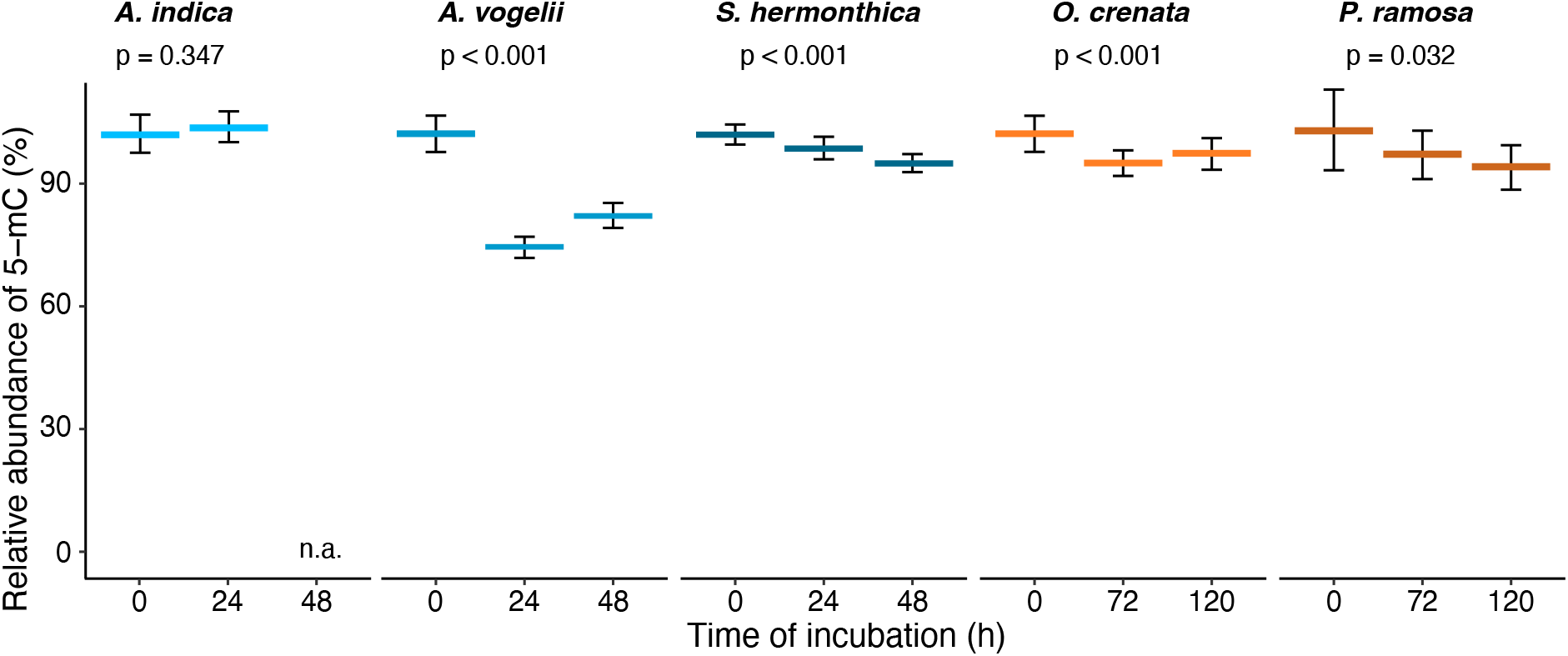
Global DNA methylation decreases along seed aging in strigolactone-dependent species. Genomic DNA was extracted from conditioned seeds that were incubated for 0−120h in a saturated sodium nitrite environment (50°C, 60% relative humidity) and used for quantifying changes in 5-methylcytosine (5-mC) abundances. Data are expressed as percentages of 5-mC relative to non-aged seeds. Data are means ± SE (n=3). *P*-values indicate the significance of the variation in 5-mC levels over seed aging (Kruskal-Wallis test).

Our data indicates that the loss of seed viability along seed aging coincides with global DNA demethylation. Similar correlations were made in non-parasitic species. For instance, a gradual decrease in germinability, viability, and 5-mC levels over three years of storage occurs in European beech (*Fagus sylvatica*) seeds, regardless the storage temperature (Michalak et al. 2023). Loss of about 50 % germinability after 18 months of cold storage of oak (*Quercus robur*) seeds was associated with a significant decrease in DNA methylation (Michalak et al. 2015b). A decrease in 5-mC levels was also demonstrated for pear (*Pyrus* communis) seeds that were cryo-stored at high water content (Plitta-Michalak et al. 2021), and decreasing 5-mC levels were detected even before any change in germination behavior in poplar (*Populus nigra*) seeds (Michalak et al. 2022). Additional lines of evidence linking DNA methylation to seed viability come from *A. thaliana* mutant lines defected in *CHROMOMETHYLASE3* (*CMT3*) and *METHYLTRANSFERASE1* (*MET1*) that displayed 23 % and 33 % of improperly developed embryos, resulting in 16.3 % and 12.2 % seed abortion, respectively (Xiao et al. 2006). Although our results align well with the literature, we expect that the extent of DNA demethylation in the context of an accelerated aging protocol differs from that of naturally aged populations. Yet, given that most differentially methylated positions within or outside gene bodies have been presupposed to be functionally inconsequential (Gallusci et al. 2023, Quadrana and Colot 2016), we assume that even minor changes in DNA methylation are sufficient to alter the function of major genes and, therefore, to exert major effect on seed viability.

## CONCLUSION & PERSPECTIVES

We showed that seed viability in strigolactone-dependent weedy broomrapes and witchweeds declines similarly as seeds age in correlation with global DNA demethylation and reduced strigolactone sensitivity. Broomrapes appear to have longer-lived and environmentally more resilient seeds than witchweeds. This may reflect adaptations to different ecological habitats (Mediterranean vs. subtropical climate) or the parasitic specialization of the investigated species (obligate hemiparasitism vs. holoparasitism). It takes only one or few individual(s) to replenish an aging seed bank. Yet, long-term persistence is a multi-layered phenomenon. First, the residual proportion of viable seeds in old seed banks are significantly less vigorous, exemplified by reduced seedling emergence or arrested development in aged, hypomethylated seeds (Michalak et al. 2023, Plitta-Michalak et al. 2022). Second, epigenetic modifications, especially those involving DNA methylation, may contribute to transgenerational epigenetic variations and priming. It remains an unanswered question in plant biology as to whether and how the transmission of epimutations over generations affects the stability of a given weed population, or if the epigenetic landscape is reset between generations. Third, the reduced sensitivity to strigolactones in aging seeds results in falling probabilities for old seed banks to re-populate a field, an effect that could be taken advantage of in parasitic plant management by aligning crop densities to seed bank ages. This study lays the groundwork for further research into identifying methylome-guided gene functions underlying parasitic plant seed responses to their environment. Future research will need to determine whether optimal environmental conditions for germination, including e.g. temperature, light quality and quantity, and soil nutrient content, vary over seed aging.

## ACKNOWLEDGMENTS

We thank the Delavault lab for sharing with us seeds of *Phelipanche ramosa*, the Runo lab for providing seeds of *Striga hermonthica* and *Alectra vogelii*, and the Aly lab for collecting for us seeds of *O. crenata*. This research received financial support from the German Science Foundation (DFG, WI 4507/3-1 to S.W.) and the German-Israeli Foundation for Scientific Research and Development (GIF, G-2415-413.13/2016 to S.W.).

## AUTHOR CONTRIBUTIONS

Designed this research: G.B. and S.W.; Conducted the experiments: G.B. and J.P.; Data analysis G.B., J.P., and S.W.; Wrote the manuscript: G.B. and S.W.

